# SpliceVisuL: Visualization of Bidirectional Long Short-term Memory Networks for Splice Junction Prediction

**DOI:** 10.1101/451906

**Authors:** Aparajita Dutta, Aman Dalmia, R Athul, Kusum Kumari Singh, Ashish Anand

## Abstract

Neural models have been able to obtain state-of-the-art performances on several genome sequence-based prediction tasks. Such models take only nucleotide sequences as input and learn relevant features on its own. However, extracting the interpretable motifs from the model remains a challenge. This work explores various existing visualization techniques in their ability to infer relevant sequence information learned by a recurrent neural network (RNN) on the task of splice junction identification. The visualization techniques have been modulated to suit the genome sequences as input. The visualizations inspect genomic regions at the level of a single nucleotide as well as a span of consecutive nucleotides. This inspection is performed based on modification of input sequences (perturbation-based) or the embedding space (back-propagation based). We infer features pertaining to both canonical and non-canonical splicing from a single neural model. Results indicate that the visualization techniques produce comparable performance for branchpoint detection. However, in case of canonical donor and acceptor junction motifs, perturbation based visualizations perform better than back-propagation based visualizations and vice-versa for non-canonical motifs.

## Introduction

The recent trend in genome sequence analysis is the application of neural network models that learn features from the sequence *de-novo*.^1,2^ The primary motivation to let the model learn relevant features by itself is to avoid the existing knowledge bias. Several deep and shallow neural network models have been successfully applied on genome sequence-based tasks, such as, identification of transcription factor binding sites,^3^ microRNA target prediction,^4^ and prediction of DNA methylation states.^5^ However, the inference of biologically relevant information learned by the models remains a challenge. In the domain of computer vision, image processing, and natural language processing (NLP), several visualization techniques have been effectively applied to analyze learned models and infer relevant features. Our work aims to apply various visualization techniques to identify the features that contribute to the prediction performance. Towards this aim, we also ask the following questions-*Q1: How can various visualization techniques be adapted for identifying the relevant features for a particular task? Q2: Do all visualization techniques deliver similar results or are one method superior to the other?*

We employ five different visualization techniques by modulating them to suit the genome sequences as input. The visualization techniques chosen can be grouped into two categories, namely intrinsic visualization and post-hoc visualization, based on the time when visualization is obtained.^6^ Intrinsic visualization has been achieved by adding an interpretable component, namely *attention layer*, to the model. This visualization technique identifies features considering all the sequences present in the dataset. The post-hoc visualization techniques employed can be further categorized into back-propagation based techniques and perturbation based techniques. This set identifies motifs based on individual sequences. The post-hoc visualization techniques used are *smooth gradient of noisy nucleotide embeddings*, *integrated gradients of nucleotide embeddings, omission of a single nucleotide*, and *occlusion of k-mers*. Among these visualization techniques, the first two are back-propagation based whereas the next two are perturbation based.

Towards our aim, we consider the task of splice junction classification and evaluate the different visualization techniques in their ability to infer relevant features known to be important for splice junction identification. Splice junction classification is an important sub-task of genome annotation. This task involves identification of exon-intron (donor site) and intron-exon (acceptor site) boundaries which are usually characterized by canonical motif dimers *GT* and *AG* respectively (further explained in Supplementary Materials (Section 1.1)). However, there are exceptions to these consensus motifs which yield the non-canonical motifs that correspond to non-canonical or unconventional splicing events.^7^ Most of the existing computational methods focus only on the identification of canonical splice junctions due to the lack of consistent non-canonical consensus. Nevertheless, the non-canonical splicing signals are equally important in understanding the splicing phenomenon, ^7^ and hence this remains an interesting area to be investigated further.

Neural network based models like^8–10^ and^11^ identify canonical and non-canonical splice junctions but with limited or no inference of learned information. Learning models like,^12,13^ and^14^ consider only the canonical splice junctions. Also, these works applied one chosen visualization technique for inference of learned features. No discussion on the comparative study of various visualization methods have been provided. This works fills these gaps by application and comparison of various visualization methods on extraction of different known canonical and non-canonical features.

Motivated by application of RNN in sequence-based bioinformatics problems,^4,10,15^ we further explore its application in splice junction prediction by employing bidirectional long short-term memory (BLSTM) units^16^ in the hidden layer. There have been earlier attempts at applying other RNN units like long short-term memory (LSTM) unit and gated recurrent unit (GRU) in this task.^10^ We further superimpose the model with an attention^17^ layer to add interpretability to the model. The contributions of this paper can be summarized as follows:

- We explore the application of BLSTM network along with attention mechanism for the prediction of splice junctions. The proposed architecture achieves the state-of-the-art performance.
- We generate two different types of negative dataset to test the consistency of recurrent neural network models compared to other neural network models.
- We redesign some of the effective visualization techniques, available in the literature, to be capable of comprehending genome sequences as inputs.
- We infer relevant biological information learned by the model for both canonical and non-canonical splicing events. The splicing features are validated with the existing knowledge from literature.
- We further provide a comprehensive study of the ability of the visualization techniques in inference of various known canonical and non-canonical features.

## Related work

### Visualization of sequence-based features

With the objective of deciphering the reason behind outstanding performance of various learning models, attempts have been made to monitor the change in model weights as learning progresses.^4,8,5^ extracts sequence motifs by aligning sequence fragments that maximally activated the filters of the convolutional layer for predicting single-cell methylation states.^18^ performs one-dimensional global average pooling on the attention weighted output of an RNN to discover the parts of the sequence that are significant for identifying pre-miRNAs.

^19^ explores various sequence-specific as well as class-specific visualization techniques to obtain the important nucleotide positions present in a genome sequence for classification of transcription factor binding sites. In contrast to their work, we have incorporated variable length occlusion to study variable length canonical as well as non-canonical splicing features apart from accessing the importance of each nucleotide position using all the visualization techniques. We also apply smooth gradient of noisy nucleotide embeddings, rather than the raw gradients, to generate sharper *sensitivity maps*.^20^

With respect to deciphering the splicing phenomenon, Zuallaert et al.^12^ apply back-propagation based visualization technique called *DeepLIFT*^21^ over a deep convolutional neural network (CNN) to infer features relevant to splicing. This model predicts either the donor or the acceptor splice sites based on the dataset used. Also, they consider only canonical splice junctions for visualizing the important genomic regions. Zhang et al.^13^ employ deep CNN for identification of canonical and semi-canonical^22^ splice junctions. They further use deep Taylor decomposition^23^ to assess the contribution of nucleotides in the classification decision. Jaganathan et al.^14^ annotate the complete RNA transcript by classifying each nucleotide as a donor, acceptor or neither. They apply dilated convolution layers to enable learning of sequence determinants from thousands of flanking nucleotides. They further extend the model to evaluate the effects of genetic mutations on splicing. They eventually enhance the understanding of the relationship between mutations in the noncoding genome and various human diseases.

### Splice junction classification

Several computational methods have been proposed for splice junction classification. In recent times, advanced sequencing technologies like RNA-seq have produced a plethora of sequenced genome. The abundance of annotated data has boosted both alignment based and machine learning based methodologies for predicting splice junctions. However, in the alignment based methods, there is a possibility of a short read randomly matching a large reference genome containing multiple occurrences of the short read sequence.^24^ Also, the existing alignment based methods^25,26^ consider only canonical splice junctions in the prediction task. ^8^

The traditional machine learning based splice junction predictors use hand-crafted features like presence or absence of specific nucleotide patterns around the splice junctions.^27,28^ Since all the splicing signals are still not known, the hand-engineered features may adversely affect the accuracy of prediction models due to the inclusion of irrelevant features as well as high dimensionality. There have been attempts at handling the issue of high dimensionality by optimizing the features using feature selection techniques.^29,30^ Nonetheless, limited biological knowledge still ensued the inclusion of irrelevant features. This revealed the necessity of applying learning techniques that can capture, by itself, the complex splicing signals from the genome sequence.

^8^ proposed a splice junction prediction model based on deep Boltzmann machine. ^9^ employed a deep CNN, named *DeepSplice* that predicts novel splice junctions.^10^ already explored different RNN units in the hidden layers of a deep neural network for predicting splice junctions but with limited exploration of model interpretability. ^11^ proposed distributed feature representations of splice junctions, named *SpliceVec*, which captures splicing features to be classified by a multilayer perceptron (MLP). These prediction models yielded promising accuracy in the prediction of splice junctions. However, most of these models fail to extract the sequence motifs that govern the splicing phenomenon due to lack of interpretability in the model.

## Methods

In this section, we introduce the neural architecture employed for classification of true and decoy splice junctions. Further, we discuss the visualization techniques applied to analyze the features learned by the model.

### Neural architecture

The overview of *SpliceVisuL* architecture is shown in Figure 1. Following is the explanation of the entire workflow.

**Figure 1:**
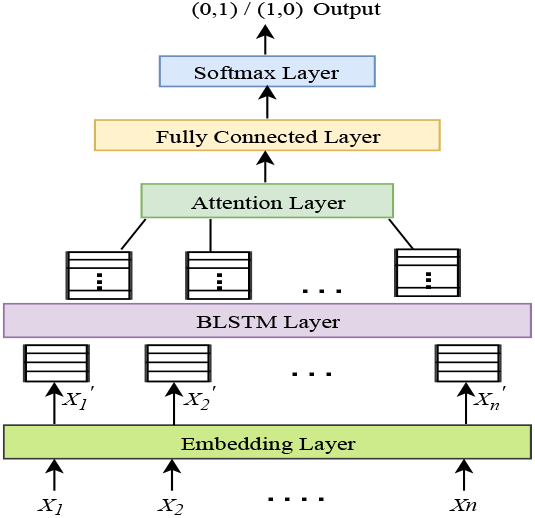
An overview of SpliceVisuL architecture.

#### Input representation

Input is a putative splice junction sequence consisting of five nucleotide codes, *A (Adenine), C (Cytosine), G (Guanine), T (Thymine) and N (representing any one of the four nucleotides*), where each of the nucleotide code is represented as an integer. We use a dense vector representation for each of the five nucleotide codes. Each input sequence is passed through an embedding layer which transforms each input splice junction sequence of length n into an *n* × 4 dimensional dense vector that gets updated while training the neural network.

#### Modeling splice junctions using BLSTM network

The *n* × 4 dimensional dense input vectors are fed in mini-batches into both the forward and backward LSTM^31^ layers configured as a BLSTM (details on LSTM and BLSTM provided in Supplementary Materials (Section 1.2)). Both the LSTM layers learn meaningful features in a supervised manner to generate an *n* × *n_l_* dimensional vector representation, where *n_l_* is the number of hidden units in each LSTM layer. Both the vectors generated by the forward and backward LSTM layers are concatenated to generate an *n* × 2*n_l_* dimensional vector representing the learned features of each splice junction.

#### Feature interpretation using attention layer

The attention layer is added to obtain a more targeted model which can capture the role each nucleotide in the input sequence plays in classification of the splice junction. This layer adds the ability of intrinsic visualization to the model. We implement the traditional attention mechanism proposed in^17^ (explained in Supplementary Materials (Section 1.2)). The *n* × 2*n_l_* representation of each sequence obtained from the BLSTM network is fed into the attention layer. The attention weights obtained from this layer are fed into a fully connected layer and eventually to a softmax layer to obtain the classification results. We use binary cross-entropy and Adam^32^ as the loss function and the optimizer respectively.

### Visualization techniques

The visualization techniques rely on measuring the change in performance of the learned model effected by changing either the input sequence or the embedding space of the model for a genomic region. The change can be implemented at a single nucleotide or a span of consecutive nucleotides. These visualization techniques add post-hoc interpretability to the model. The post-hoc visualization techniques used in this work can be categorized as perturbation based and back-propagation based visualizations.

Perturbation based visualization techniques used are based on modification/masking of the input sequence. Back-propagation based techniques used are based on modification of the embedding space of the model. In some sense, the perturbation based techniques mimic the site-specific mutagenesis^33^ performed in a wet lab setup. The visualization techniques are applied to define a scoring function, referred to as the *deviation value*. The *deviation value* of a genomic region reflects the contribution of that region to the classification score. We evaluate the visualization techniques, with respect to the inference of known splicing features, based on these *deviation values*. The various visualization techniques employed are described in the following subsections.

#### Smooth gradient with noisy nucleotide embeddings

In image classification tasks, the gradient of the unnormalized output probabilities with respect to the input image, referred to as *sensitivity maps*,^20^ indicate how much can a tiny change in that pixel affect the final output. Thus, to incur a minute change in the input sequence, we add noise to the embeddings of the nucleotides and compute the change in classification score. However, the sensitivity maps resulting from raw gradients are usually noisy.^34^ Therefore, based on the concept of smooth gradient,^34^ we average out the gradients obtained from several different noisy embeddings for each position of a sequence. For each input sequence, we generate 50 samples of embeddings by adding Gaussian noise with a standard deviation of 0.15. The average gradient at each sequence position, named *smooth gradient*, is the *deviation value* in this case. As a result, one might expect that the resulting averaged sensitivity map would crisply highlight the key regions.

#### Integrated gradient with nucleotide embeddings

Similar to the smooth gradient with noisy nucleotide embeddings, integrated gradient is also based on the computation of sensitivity maps. However, instead of computing a single gradient of the output probability with respect to the input, integrated gradient computes the average gradient as the input varies stepwise along a linear path from a baseline to the given input.^35^ We consider 50 steps for our experiments. The average gradient at each sequence position is the *deviation value* here. The baseline in our case is the zero embedding vector as suggested in^35^ for text inputs. We choose this visualization technique over other gradient based visualizations like DeepLIFT as it satisfies two desirable properties for attribution methods: *sensitivity* and *implementation invariance*.^35^ Also, DeepLIFT diverges from integrated gradients and does not produce meaningful results when applied to RNNs with multiplicative interactions (LSTM and BLSTM units) as it does not satisfy the *completeness* property in that case.^36^

#### Omission of a single nucleotide

The feature vector obtained from the fully connected layer of *SpliceVisuL* represents the complete input sequence. To measure the significance of each sequence position, we calculate its *omission score*.^37^ The omission score of the *j^th^* position 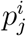 in a sequence *s_i_* is given by

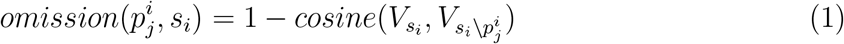

where *V_s_i__* is the feature representation obtained from the fully connected layer for the sequence *s_i_* and 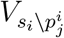 is the feature representation obtained from the same layer for the same sequence with the nucleotide at position 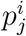 replaced by 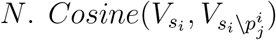 measures the similarity of the two vectors and is calculated as

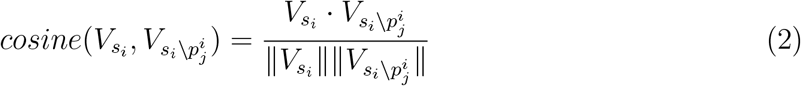

Therefore, omission score measures the *deviation value* of the vector representations of the sequence, with and without the omitted nucleotide. Higher deviation implies a higher significance of that sequence position.

#### Occlusion of k-mers

We occluded portions of a sequence to observe the variation in the predicted output. This approach has its motivation from.^38^ For each sequence, we run a sliding window *w_l_* of length *l*, centered at nucleotide number (*l* + 1)/2, and replace the nucleotides within the window with *N*. We pass the modified sequence through the model to obtain *deviation value* given by the absolute difference of model outputs with and without occlusion. For occlusion of a window of length *l*, the *deviation value* is stored in the center, that is, in position (*l* + 1)/2, of the window.

We generate *deviation values* for test sequences in batches. For a batch of size *B*, the *deviation values* are in the form of a matrix of size *B* × *n* where each input sequence is of length *n*. The computationally efficient implementation is explained in Supplementary Materials (Section 2.1). We propose two variations of occlusion described as follows:

##### Fixed length occlusion

This considers occlusion of k nucleotides (denoted by *occlusion-*) where *k* is 1 or 3 in our experiments. *Deviation values* at boundary indices are computed by occluding the first and last (*k* + 1)/2 indices. The significance of a genomic region is proportional to the corresponding *deviation value*.

##### Variable length occlusion

Fixed length occlusion has a limitation of considering fixed length genomic regions whereas in real scenario there may be sequence patterns of variable lengths that regulate splicing. Hence, we incorporate variable length occlusion where for each index *j* of a sequence *s_i_*, we occlude a window *w_l_* of length *l* ∈ {1, 3, 5, 7, 9, 11}. We compute the *deviation values* denoted by 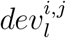, for each window length *l*, with and without occlusion. The *deviation value* assigned to position (*l* + 1)/2 of window *w_l_* is given by 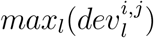 for *l* ∈ {1, 3, 5, 7, 9, 11}. The window length 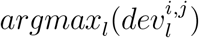 corresponding to index *j* of sequence *s_i_* is stored in the *j^th^* column of the *i^th^* row of a *window matrix*. Therefore, the value in *j^th^* column of *i^th^* row of the *window matrix* signifies the length of the pattern, centered at index *j* of sequence *s_i_*, that contributes maximum to the prediction of the model.

### Prerequisites for visualization

For each of the visualization techniques, we extract the relevant features learned by the model using the following methods over the sets of canonical and non-canonical test sequences separately. Going forward, the splice junction sequences comprising the canonical dimer motifs (*GT-AG*) are referred to as the canonical sequences and likewise for the non-canonical sequences. The first three methods are adapted from^12^ for analyzing the same splicing features as discussed in,^12^ but considering both canonical and non-canonical sequences.

1. Average deviation value per position: At each sequence position, we compute the average absolute deviation values across all the sequences irrespective of the nucleotide type
2. Average deviation value per position per nucleotide: At each sequence position, we compute the average deviation values per nucleotide across all the sequences.
3. Average deviation value of a specific pattern in a specific region of the sequence: We compute the deviation value of a specific pattern by summing up the deviation value of each nucleotide in the pattern. For each starting position in the specific region, we calculate the average deviation values of all occurrences of the pattern across the sequences.
4. Frequency of different occlusion windows per position: We compute the frequency of different window lengths per position based on the number of times occlusion of that window length centered at that position contributed to maximum deviation values across all sequences. This means that we compute the frequency of window lengths per position based on the *window matrix* explained in previous subsection.

## Experimental setup

### Positive data generation

We use GENCODE annotations,^39^ based on human genome assembly version *GRCh38*, to train and test our model. We target to assess the model’s performance on the prediction of novel splice junctions. Several alignment based methods^25,26^ have the potential to identify novel splice junctions through *ab initio* alignment of RNA-seq reads to the reference genome. Machine learning based approach ^9,13^ was also applied recently for identification of novel splice junctions. To this end, we train the model using an earlier release (version 20) and test the model on only the newly added splice junctions in a later release (version 26), as described in.^9^

Unlike some contemporary state-of-the-art models like^8,12^ that consider either acceptor or donor splice sites as the input data, we consider junction pairs as the input for training and testing the model as in.^9,11,13^ This results in performance improvement in the model as shown in Table 1 of Supplementary Materials (Section 2.2). This can also be justified by the study that suggests that splice sites are not recognized independently through individual consensus but usually in pairs across exons or introns. ^40,41^

**Table 1:** Performance of *SpliceVisuL* compared with state-of-the-art models. Accuracy (Ac), Precision (Pr), Recall (Re) and F1 Score (F1) are computed in percentage.

| Model | Type-1 Dataset | Type-2 Dataset |
| --- | --- | --- |
|  | *Ac, Pr, Re, F1*(%) | *Ac, Pr, Re, F1*(%) |
| SpliceVisuL | 98.24, 99.66, 96.81, 98.21 | 98.38, 99.38, 97.38, 98.37 |
| LSTM | 98.19, 99.45, 96.91, 98.16 | 98.36, 99.38, 97.34, 98.35 |
| LSTM with attention | 98.24, 99.70, 96.77, 98.21 | 97.68, 98.88, 96.45, 97.65 |
| SpliceRover | 96.03, 95.98, 96.10, 96.04 | 90.13, 89.90, 90.41, 90.16 |
| DeepSplice | 89.81, 94.77, 84.43, 89.05 | 92.85, 96.43, 89.03, 92.56 |
| SpliceVec-MLP | 71.57, 71.42, 72.28, 71.72 | 93.98, 96.43, 91.40, 93.78 |

We extract 291,831 and 294,576 unique splice junction pairs from version 20 and version 26 respectively. The junction pairs are extracted from protein-coding genes only. Each junction pair comprises an intron with flanking upstream and downstream exonic regions. We consider introns of length greater than 30 base pairs (bp) based on a study^42^ which states that introns shorter than 30 bp usually result from sequencing errors in the genome.^11^ This reduces the number of junction pairs to 290,502 and 293,889 in versions 20 and 26 respectively. Our test data comprises 5,612 novel junction pairs present only in version 26.

### Negative data generation

Existing works usually adopt one of the following two techniques to generate negative data. We adopt both the ways of generating the negative data to have comprehensive and unbiased analysis. The Type-1 dataset is generated similar to the standard procedure described in.^43^ We extract a portion of the sequence from the center of each intron to generate a pseudo sequence. The length of the extracted portion is kept equal to the length of an input sequence. This set of negative data captures the non-randomness of DNA sequences. We obtain 290,502 false samples for training data and 5,612 false samples for testing data using this procedure.

More than 98% of splice junctions contain the consensus dimer GT and AG at the donor and acceptor junctions respectively.^22^ However, the occurrence of the consensus dinucleotide is far more frequent compared to the number of true splice junctions in the genome. The neat exclusion of the pseudo sites by the splicing mechanism suggests the presence of other subtle splicing signals which play an important role in the process. The presence of consensus dimer in all the negative samples will result in the model learning the remaining splicing patterns present in the vicinity of the splice junctions.

Therefore, we generate the Type-2 dataset based on the procedure described in^9,11^ where the negative data is randomly sampled from the human genome assembly version *GRCh38*. For each decoy junction pair, we randomly search for the consensus dimer GT and AG such that both lie in the same chromosome and the distance between them lies in the range of 30 and 1,240,200 nucleotides (nt). We obtain a huge number of such samples using this procedure, out of which we randomly select 290,502 false samples for training data and 5,612 false samples for testing data. Both the scenarios are pictorially depicted in Figure 4 of Supplementary Materials (Section 2.3).

### Training and Hyperparameter tuning

Each input splice junction is truncated to 40 nt upstream and downstream flanking regions of the consensus dimer GT or AG, thus obtaining an 82 nt sequence. Both the donor and acceptor junctions of a junction pair are concatenated to form a 164 nt sequence. The effect of variation in the flanking region on model accuracy is shown in Table 2 of Supplementary Materials (Section 2.4).

**Table 2:** Summary of the various visualization techniques in their ability to identify selected canonical splicing features.

| Features | Attention | Occlusion | Omission | Smooth gradients | Integrated gradients |
| --- | --- | --- | --- | --- | --- |
| Importance of sequence position | ✓ | ✓ | ✓ | ✓ | ✓ |
| Donor site consensus | ✗ | ✓ | ✓ | ✗ | ✓ |
| Acceptor site consensus | ✗ | ✓ | ✓ | ✓ | ✗ |
| Branchpoint | ✓ | ✓ | ✓ | ✓ | ✓ |

*SpliceVisuL* can be represented as a (1-4-100-100-2048-2) architecture where 4-dimensional embeddings are passed through a BLSTM, attention and fully connected layer with 100, 100 and 2048 units respectively. Values for batch size, dropout, recurrent dropout, and epochs are tuned to 128, 0.5, 0.2 and 50 respectively. The hyperparameters are tuned by partitioning the training data from version 20 into 90% training and 10% validation data. All experiments were carried out on an NVIDIA GeForce GTX 980 Ti GPU machine with 6GB memory. We evaluate the performance of the classifier based on precision, recall, accuracy, and F1 score. Table 3 in Supplementary Materials (Section 2.5) shows the variation in the performance of the model with variation in the number of hidden layers.

### Baselines

We implemented the following state-of-the-art models as baselines and compared the results obtained by the various models on the same set of training and testing data. The hyperparameters for the baselines (details in Supplementary Materials (Section 2.6)) were tuned using the same process mentioned in previous subsection.

1. Vanilla LSTM: This model comprises an embedding layer, two hidden LSTM layers, and a softmax output layer as proposed in.^10^
2. LSTM with attention: We replaced the hidden units of the proposed architecture with LSTM units.
3. SpliceRover: This model classifies an input sequence as donor (acceptor) or not donor (acceptor) in the donor (acceptor) classification model. The classifier comprises of multiple alternating convolutional and max-pooling layers.^12^
4. DeepSplice: This model classifies input sequences using a deep CNN comprising of two convolutional layers.^9^ Input sequences are represented in the form of a 4 × *N* matrix comprising the flanking ‘N’ nucleotides in the upstream and downstream of both acceptor and donor splice junctions.
5. SpliceVec-MLP: This model learns feature vectors of the entire intronic region along with flanking upstream and downstream exonic regions to be classified using an MLP.^11^ The input formation for SpliceVec-MLP is described in Supplementary Materials (Section 2.7).

## Results

### Prediction performance

To assess the quality of feature embedding obtained from the dense layer of *SpliceVisuL*, we plot the 2048-dimensional vector by reducing it to a two-dimensional vector using Stochastic Neighbor Embedding (t-SNE).^44^ We observe that the projected representations are distinctly separable in the two-dimensional feature space. Figure 2 shows a t-SNE plot of randomly selected 1000 true and 1000 decoy splice junction pairs. Figure 6 in Supplementary Materials (Section 3.1) shows four different plots of randomly selected 1000 true and 1000 decoy splice junction pairs ^1^.

**Figure 2:**
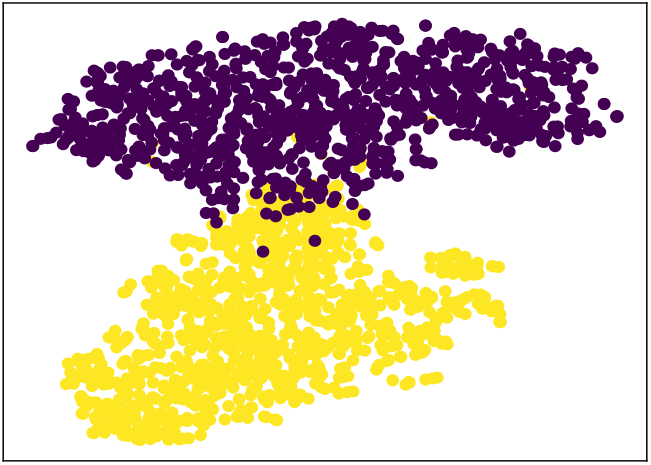
t-SNE plot of 1000 true and 1000 decoy splice junctions. Points in blue represent true whereas points in yellow represent decoy splice junction pairs.

We evaluate the performance of *SpliceVisuL* using both Type-1 and Type-2 dataset. We compare the predictive performance with the baselines discussed earlier. Table 1 shows comparison of *SpliceVisuL* with the baselines. We observe a performance improvement of the RNN based architectures, over other neural network based baselines, in the range of 2%-27% for the Type-1 dataset and an improvement in the range of 4%-8% for the Type-2 dataset. The RNN based models are consistent in its performance with both the dataset. We analyze the performance of *SpliceVisuL* on the Type-2 dataset.

### Visualization of splicing features

We apply the various visualization techniques on *SpliceVisuL* to decipher the features learned by the model for splice junction identification. The following analysis has been done on 10853 canonical and 371 non-canonical test sequences. Additional visualizations are shown in Supplementary Materials (Section 3).

#### Identifying the significance of sequence positions near splice junctions

All the visualization techniques identify nucleotides in the proximity of splice junctions as the most significant based on average deviation value per position. The nucleotide importance decreases as we move further in the flanking region. Figures 3(a) (3(c)) and 3(b) (3(d)) show the deviation values obtained from occlusion-1 (smooth gradients) at canonical donor and acceptor splice junctions respectively. Figures 3(e) and 3(f) show the deviation values obtained from occlusion-1 and smooth gradients at non-canonical acceptor splice junctions respectively.

**Figure 3:**
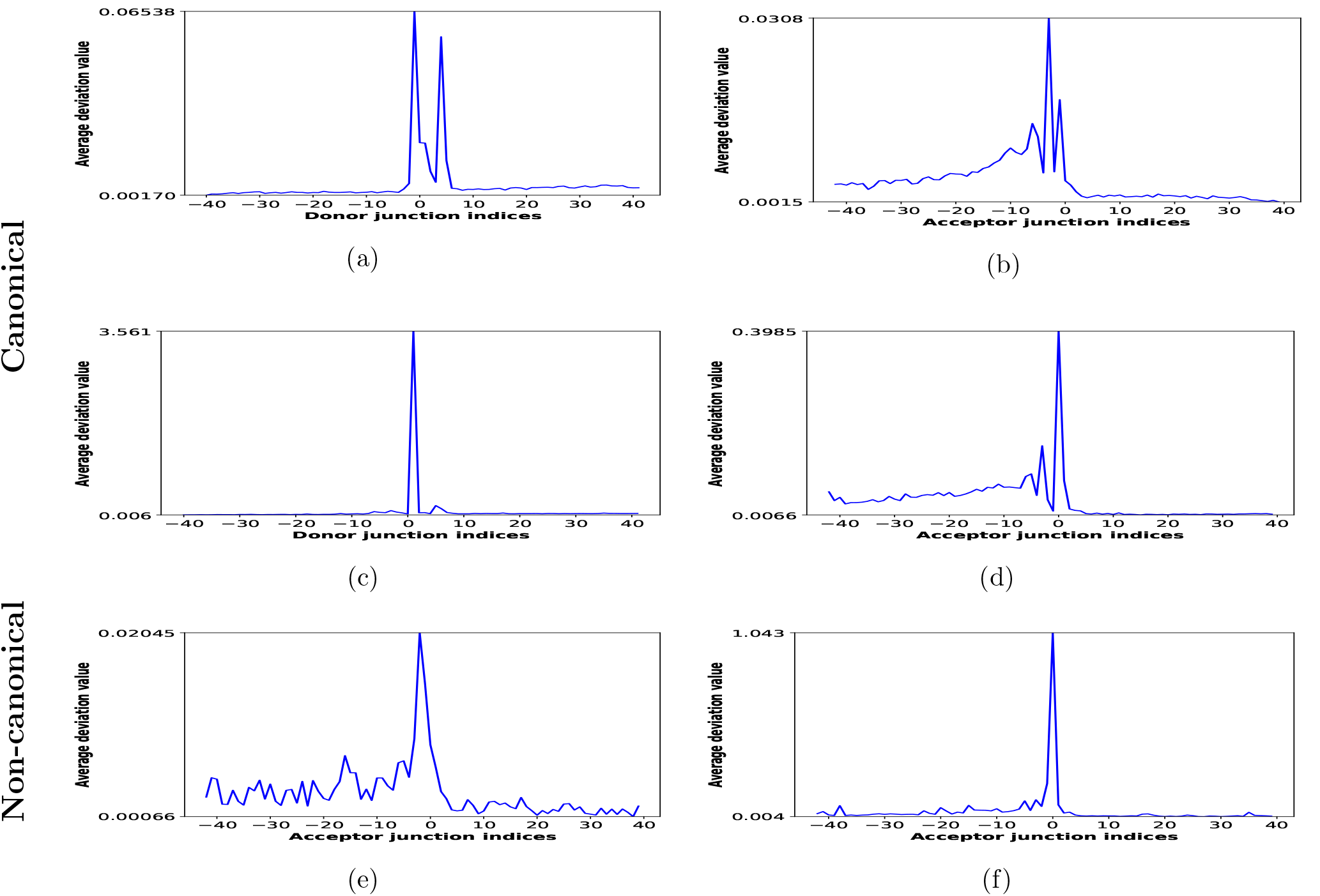
Significance of sequence positions near splice junctions. The average deviation value per nucleotide position is shown for occlusion-1 of canonical (a) donor and (b) acceptor splice junctions, smooth gradient of canonical (c) donor and (d) acceptor splice junctions, (e) occlusion-1 of non-canonical acceptor and (f) smooth gradient of non-canonical acceptor splice junctions.

**Figure 4:**
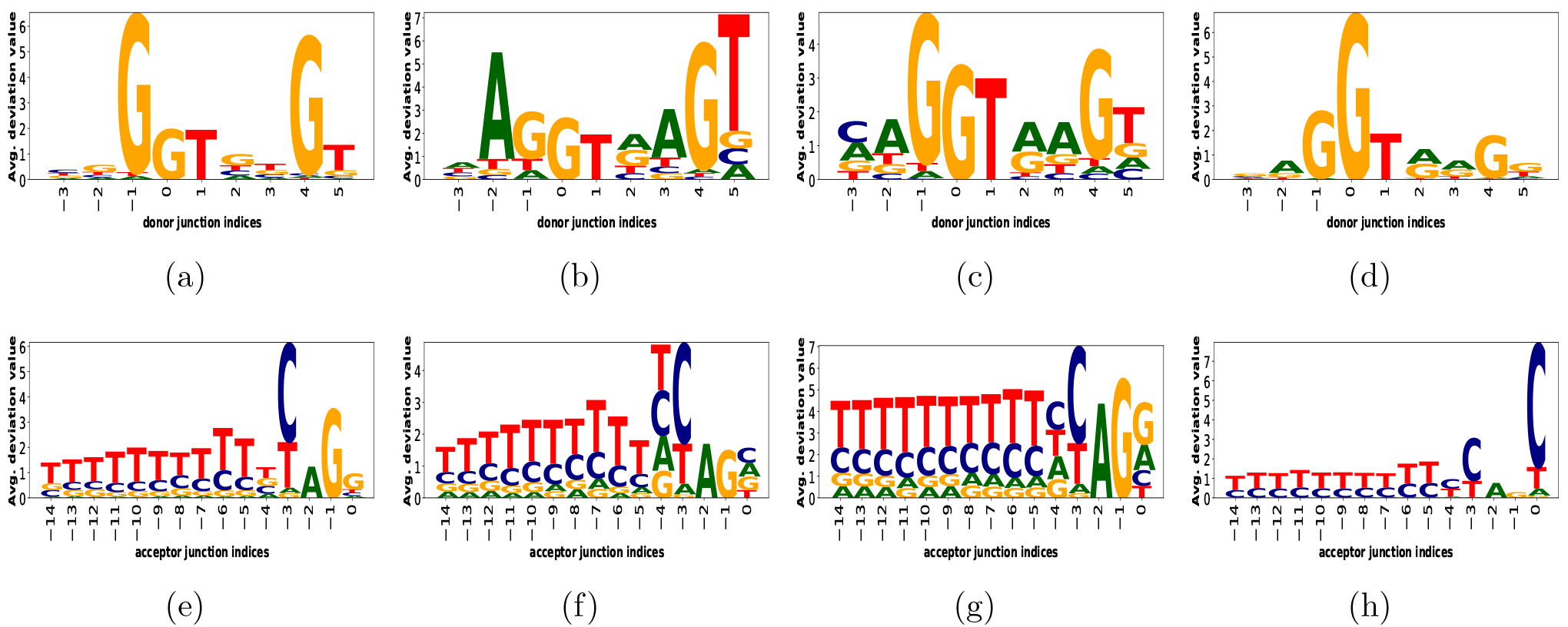
The splicing motifs at canonical splice junctions. The average deviation value per position per nucleotide is shown for canonical donor junction motifs obtained from (a) occlusion-1 (b) occlusion-3 (c) omission (d) integrated gradients and canonical acceptor junction motifs obtained from (e) occlusion-1 (f) occlusion-3 (g) omission and (h) smooth gradients.

We observe a relatively higher importance of the intronic region at canonical acceptor splice junction due to the presence of polypyrimidine tract (PY-tract, see next subsection). The importance assigned is similar for both perturbation based occlusion and back-propagation based smooth gradients in case of canonical splice junctions.

However, in case of non-canonical splice junctions, both these techniques show different trends. Smooth gradients assign relatively lower importance to the PY-tract which is consistent with the knowledge that non-canonical splice junctions have weakly conserved PY-tract.^7^ Occlusion, on the other hand, assigns relatively higher importance to the upstream region of the acceptor junction.

The higher importance assigned by occlusion can be attributed to a characteristics of perturbation based visualization techniques explained in. ^36^ The characteristics illustrates that back-propagation based methods are better in capturing the effect of multiple features together whereas perturbation based methods are better in explaining the role of individual features in isolation. Considering each nucleotide position as a feature, occlusion assigning higher importance to the upstream region suggests that features in this region work in isolation, rather than as consecutive consensus, in case of non-canonical splicing.

#### Identifying the splicing motifs at splice junctions

##### Canonical motifs

We compare the obtained splicing motifs with the existing consensus of canonical splice junctions known from literature. Figure 5 in Supplementary Materials (Section 2.8) shows the extended donor site consensus 9-mer [*AC*]*AGGTRAGT* and the extended acceptor site consensus 15-mer *Y*_10_*NCAGG*, known from existing studies^45^ and also complies with that in our dataset. The acceptor site consensus mostly consists of the PY-tract (*Y*_10_).

**Figure 5:**
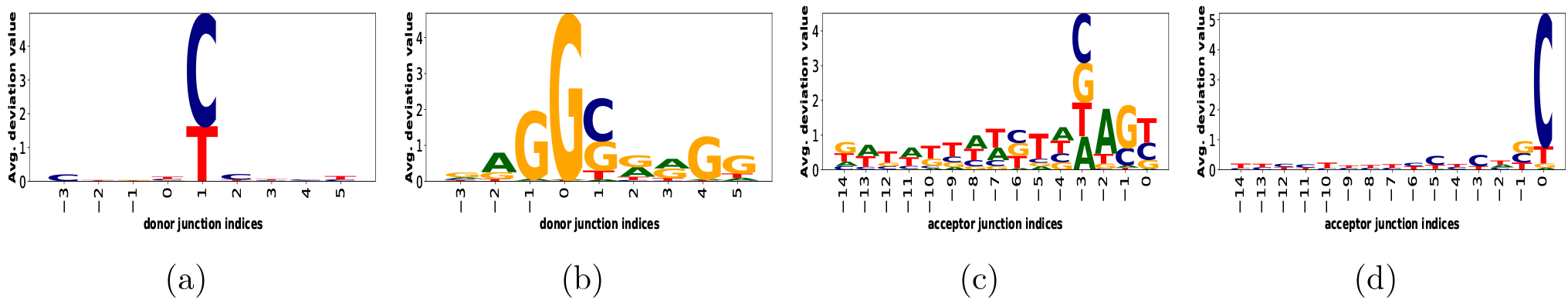
The splicing motifs at non-canonical splice junctions. The average deviation value per position per nucleotide is shown for non-canonical donor junction motifs obtained from (a) smooth gradients and (b) integrated gradients as well as non-canonical acceptor junction motifs obtained from (c) occlusion-3 and (d) smooth gradients.

Figures 4(a), 4(e), 4(c) and 4(g) show the donor and acceptor junction motifs captured by occlusion-1 and omission respectively. Both the visualizations capture most of the known consensus. We also observe that the donor and acceptor junction motifs (Figures 4(b) and 4(f)) captured by occlusion-3 are better than occlusion-1. This is seen as the inherent behavior of occlusion that larger occlusion windows imply better representations learned.^36^ This is also observed in our results obtained for identification of branchpoint.

However, back-propagation based visualizations capture motifs partially. Integrated gradients learn better representation for donor junction (Figure 4(d)) whereas smooth gradients learn better representation for acceptor junction (Figure 4(h)). However, a back-propagation based visualization technique named DeepLIFT (^21^) identified both donor and acceptor junction consensus in. ^12^ This can be attributed to the factors that our dataset comprises donor-acceptor junctions pairs as well as both canonical and non-canonical sequences.

Figure 17 in Supplementary Materials shows the donor and acceptor splice junction motifs captured by attention. Attention visualization is limited in a way that it identifies importance considering overall dataset and hence does not provide information regarding what class the feature is important for.^46^ Therefore, it possibly overlooks relevant signals regulating splicing.

##### Non-canonical motifs

Two existing knowledge from non-canonical splicing is observed in the results obtained from these visualizations. Both the back-propagation based visualizations (smooth gradients (Figure 5(a)) and integrated gradients (Figure 5(b))) pick up GC as an alternative donor site consensus. This has also been observed in previous study where the most common class of non-consensus donor splice junction has been reported as *GC*. ^47^

Both occlusion-3 (Figure 5(c)) and smooth gradients (Figure 5(d)) suggest the presence of weaker PY-tract upstream of the non-canonical acceptor junction. This is also in coherence with the observation made in identifying the significance of sequence positions. Additionally, both these methods suggest a weaker presence of nucleotide *C* at consensus dimer position – 1. This complies with the second class of exception (*AT* – *AC*) to the consensus dimers reported in. ^47^

#### Identifying the location of branchpoint consensus ‘CTRAY’

We plot the average deviation value of the branchpoint pattern ‘CTRAY’^7^ in −40 to −15 nt upstream region of the acceptor splice junctions which comes adjacent to the extended 15-mer acceptor site consensus. Although the branchpoint motif is highly degenerate, it is usually observed to be strongly conserved towards the last 50 nt of the introns with a peak around −23 nt relative to the acceptor splice junction. ^48^ This is also observed in most of the visualization results (Figure 6(a), 6(b), 6(c), 6(d)) where the peak mostly lies between −25 to −20 nt.

**Figure 6:**
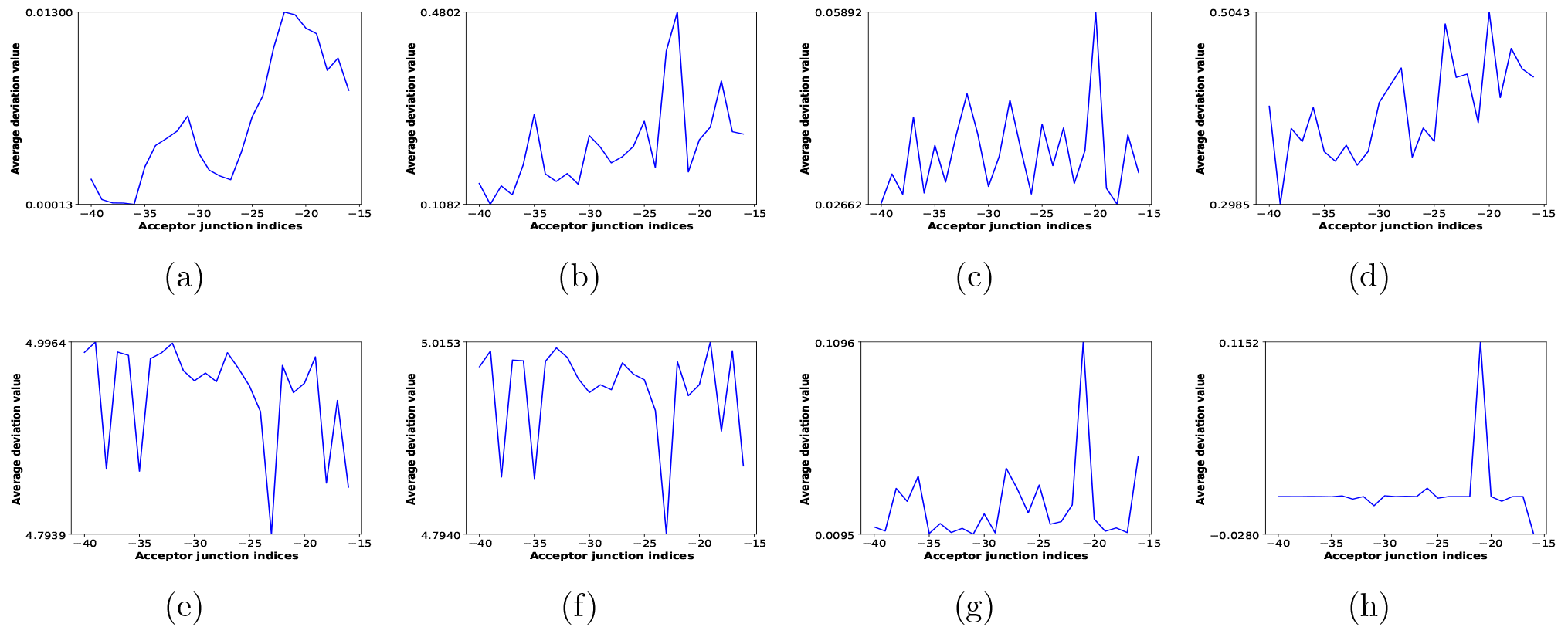
The average deviation value of branchpoint pattern CTRAY in the upstream region [−40, −15] of acceptor splice junction. The average deviation value per position for the pattern CTRAY is shown for (a) attention (b) smooth gradients (c) integrated gradients (d) omission (e) occlusion-1 (f) occlusion-3 of canonical acceptor splice junctions (g) integrated gradients and (h) occlusion-3 of non-canonical acceptor splice junctions.

The only exception occurs in case of occlusion-1 (Figure 6(e)) where peaks are observed close to −30 and −40 nt upstream. On the other hand, occlusion-3 (Figure 6(f)) displays the peak near −20 nt. This complies with the fact that in non-linear models like deep neural networks, the result of occlusion is strongly influenced by the number of features occluded where the key features are focused when using bigger occlusion windows.^36^ Branchpoint peak is observed within similar range (−25 to −20 nt) in non-canonical acceptor splice junctions as well, shown for integrated gradients (Figure 6(g)) and occlusion-3 (Figure 6(h)).

#### Identifying the optimal motif length per position

To leverage the capability of occlusion technique to focus on variable length features, we plot the frequency of different window lengths (varying from 1 to 11) with highest deviation values centered per position. We observe that for canonical donor junction (Figure 7(a)), both the dimer indices (0, 1) produce the maximum deviation values for window length 1. This suggests that these two positions alone are maximum functional in identification of donor splice junction. As we move further in the flanking region, the optimal feature length increases up to 9 suggesting that the flanking nucleotides are weaker signals that depend on the extended 9-mer motif for recognition of the splice junctions.

**Figure 7:**
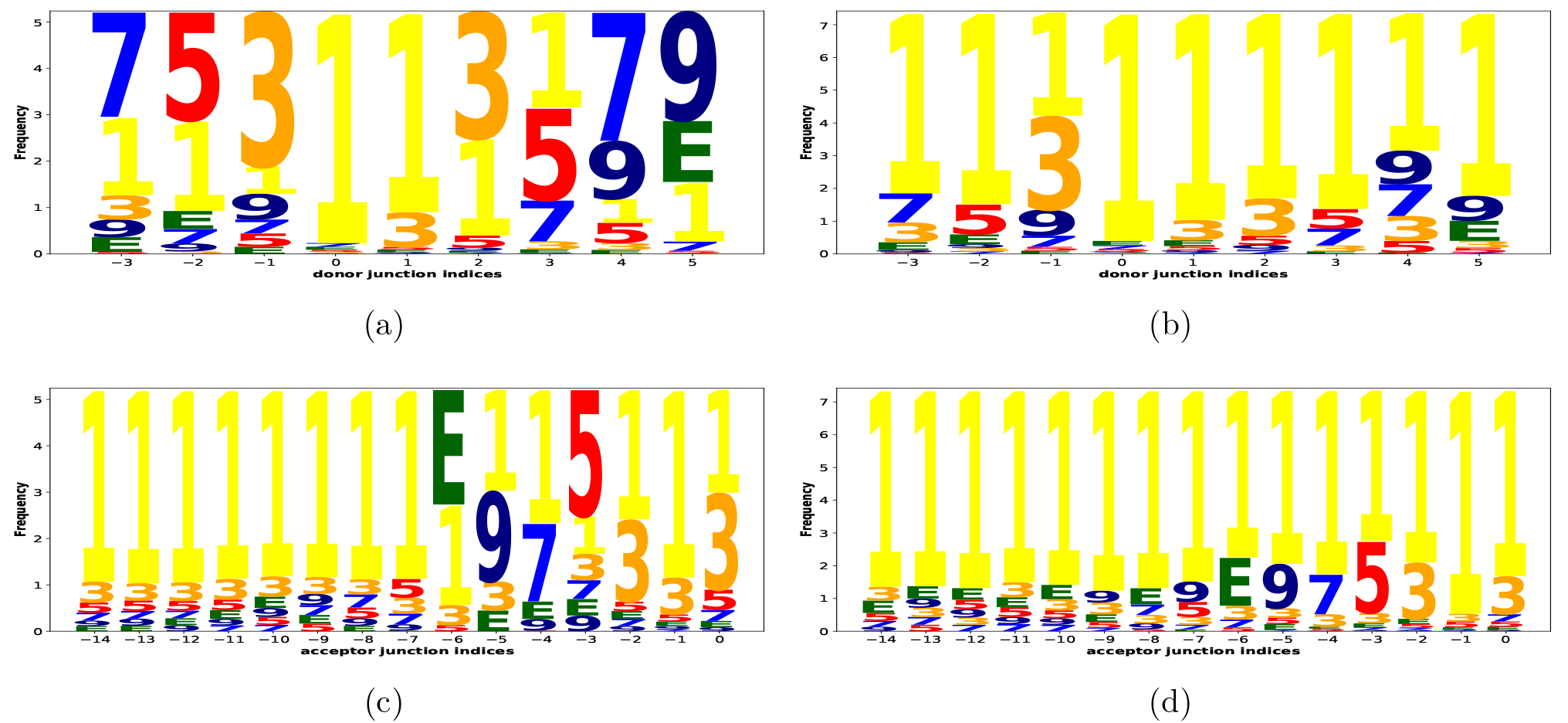
The optimal motif length per position. The frequency of different window lengths, varying from 1 to 11 (represented by E), is shown for occlusion of canonical (a) donor and (c) acceptor splice junctions and occlusion of non-canonical (b) donor and (d) acceptor splice junctions.

The canonical acceptor splice junction (Figure 7(c)) shows similar trend where optimal feature length goes upto 11 nt (denoted by ‘E’) in the PY-tract region. But, the PY-tract majorly displays 1 nt as the optimal feature length suggesting the PY-tract as another key feature for acceptor junction identification. ^7^

On the contrary, non-canonical donor and acceptor junctions (Figure 7(b) and 7(d)) consistently focus 1 nt as the optimal feature length across the junction region. This suggests that the non-canonical junction identification does not primarily depend on consecutive nucleotides in the vicinity of splice junctions but possibly to different individual nucleotides in that region. This complies with the observation in identifying the significance of sequence positions. This can also be justified by the study which shows that non-canonical splice junctions usually lack one or more known canonical consensus or the consensus may be distally located resulting in the existence of novel recognition pathways for non-canonical splicing. ^7^

## Discussion

We have done a comprehensive analysis of various visualization techniques based on the extraction of several known splicing features. The observations made on the ability of the visualization techniques, in case of canonical sequences, are summarized in Table 2. Although the visualizations display comparable performances in most of the selected features, we still conclude that perturbation based visualizations (occlusion and omission) are the most consistent in their performance across all known motifs in canonical sequences.

However, in case of non-canonical sequences, we observe a mixed performance of perturbation and back-propagation based visualizations in capturing the most commonly occurring non-canonical consensus known in literature. The understanding developed in this work regarding non-canonical splicing is far from complete.

We have the existing knowledge that non-canonical splicing signals may partially lack the known consensus or the signals may be scattered far from the splice junctions. ^7^ We also observe that non-canonical splicing signals may be in the form of various nucleotides dispersed across the genome rather than consecutive nucleotides. These results suggest the presence of non-canonical splicing signals far beyond the 40 nt flanking region considered in this work. Therefore, it would be interesting to explore larger context, especially in non-canonical sequences, to develop a better understanding of the splicing phenomenon as a whole.

## Conclusion

In this work, we achieve state-of-the-art performance by application of BLSTM network along with attention mechanism for the prediction of splice junctions. We redesign some of the existing visualization techniques to be capable of comprehending genome sequences as input for inferring biological features relevant to canonical and non-canonical splicing. We validate the splicing motifs inferred from canonical and non-canonical sequences by comparing it with the consensus known from literature. We also summarize the performances of various visualization techniques on selected splicing features.

## Footnotes

1 We plotted 1000 junctions considering the execution time required for larger sample sizes.

