## Supplementary Materials for "SpliceVisuL: Visualization of Bidirectional Long Short-term Memory Networks for Splice Junction Prediction"

#### 1 Background

##### 1.1 Biological preliminaries

Genes, in eukaryotes, comprise of alternating regions called *exons* and *introns*. During transcription, a process called splicing removes the introns and ligates the exons with the help of a complex molecular machine called *spliceosome*. Splicing occurs at exon-intron (*donor site*) and intron-exon (*acceptor site*) boundaries called *splice sites* or *splice junctions*.

However, more than 90% of multi-exon genes undergo *alternative splicing* (AS) wherein an intron may be retained and an exon may be skipped in a pre-mRNA transcript during splicing (Lu *et al.*, 2012). AS results in the formation of multiple protein isoforms from a single gene, thereby increasing the informational diversity and functional capacity of a gene. Accurate identification of splice junctions is, therefore, important for unfolding the gene structure and expression along with providing meaningful insights into its role in alternative splicing.

The spliceosome, during splicing, identifies the splice sites based on splicing consensus residing in the vicinity of the splice junctions. Splice junctions generally comprise the canonical splicing pattern GT and AG at the donor and acceptor sites respectively. Apart from this, the consensus sequence comprises an extended donor site consensus 9-mer $[AC]AGGTRAGT$ and an extended acceptor site consensus 15-mer $Y_{10}NCAGG$ containing a polypyrimidine tract (PY-tract) preceding the acceptor site (Zhang *et al.*, 2003). There is also a very weak branch site consensus $CTRAY$ located approximately 30 nucleotides (nt) upstream from the acceptor site. However, there are evidences of existence of non-canonical splice junctions that do not comply with the above consensus.

##### 1.2 Preliminaries on model architecture

**Recurrent neural networks (RNN)** An RNN is a class of artificial neural network that comprises of a hidden feedback unit which gets repeated each time an input is fed into the network. For a given input sequence $x = (x_1, x_2, ..., x_T)$, any $x_t$ where $t \in 1, 2, ..., T$, can be considered as a time step. An RNN can be unrolled along time steps, as shown in Figure 1, to form a directed graph along the input sequence. This allows it to exhibit dynamic temporal behavior for a sequence.

The weights $W_{xh}$, $W_{hh}$ and $W_{hy}$, of the input, hidden and output layers respectively, are the same across every time step. This suggests that an RNN performs the same task on every element of an input sequence, with dependency on the previous time step. The hidden state $h_t$ at time step $t$ of an RNN behaves as the memory of the network that is computed based on previous hidden state $h_{t-1}$ and current input state $x_t$. This can be represented by the following recurrence formula:

$$h_t = f(h_{t-1}, x_t)$$

where $f$ is any non-linearity function of the hidden layer. This can be further expanded as:

$$h_t = f(W_{hh}h_{t-1} + W_{xh}x_t)$$

The output at time step $t$ is computed by:

$$y_t = W_{hy}h_t$$

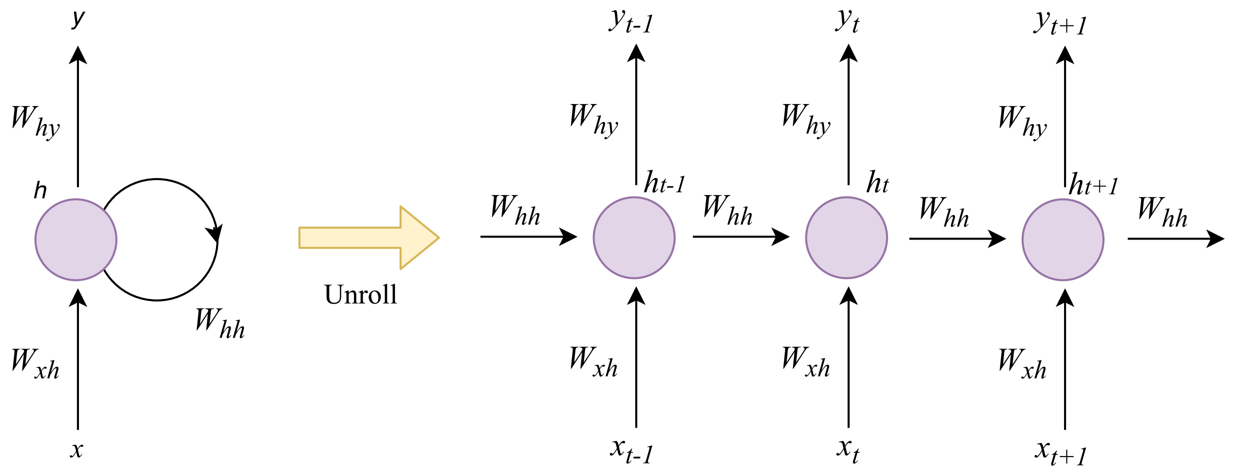


Figure 1: An unrolled RNN.

**Long short-term memory (LSTM) units** Training a vanilla RNN involves updating the input, hidden and output weights, using backpropagation through time, to reduce the total error of the network. Weight of each time step of an unrolled RNN is updated in proportion to the derivative of the error with respect to the weight. As we move along the time steps, the backward flow of the gradient results in either exploding or diminishing of the gradient exponentially. This occurs because a portion of the weight matrix gets multiplied by itself at each step of backpropagation. Hence, the vanilla RNN architecture was not capable of accessing long-term dependencies.

LSTM units are a type of building blocks, for the layers of an RNN, that are capable of memorizing long-term dependencies. LSTM units behave like memory cells comprising of input, output and forget gates. Figure 2 illustrates a single LSTM unit. The backpropagation passes through only the cell state which is controlled by the three gates. The activation vectors $i_t$, $o_t$, $f_t$, and $c'_t$, for the input gate, output gate, forget gate and candidate cell state at time step $t$, can be represented by the following equations:

$$i_t = \sigma(W_{hi}h_{t-1} + W_{xi}x_t)$$

$$o_t = \sigma(W_{ho}h_{t-1} + W_{xo}x_t)$$

$$f_t = \sigma(W_{hf}h_{t-1} + W_{xf}x_t)$$

$$c'_t = tanh(W_{hc}h_{t-1} + W_{xc}x_t)$$

The actual cell state value ($C_t$) at time step $t$ is computed by:

$$C_t = f_tC_{t-1} + i_tc'_t$$

Finally, the output ($h_t$) of the unit at time step $t$ is given by:

$$h_t = o_ttanh(C_t)$$

**Bidirectional long short-term memory (BLSTM) networks** A limitation of traditional RNN is that it makes decisions based on previous context only. However, language translation and speech recognition systems, where whole input is transcribed at once, exploiting future context is expected to improve predictive performance.

A BLSTM network is the bidirectional variant of an RNN that employs an LSTM unit in the hidden layer. Figure 3 shows a BLSTM network. It comprises of two identical hidden LSTM layers. One of the layers is fed with the input as-is whereas the other layer is fed with the reversed copy of the same input.

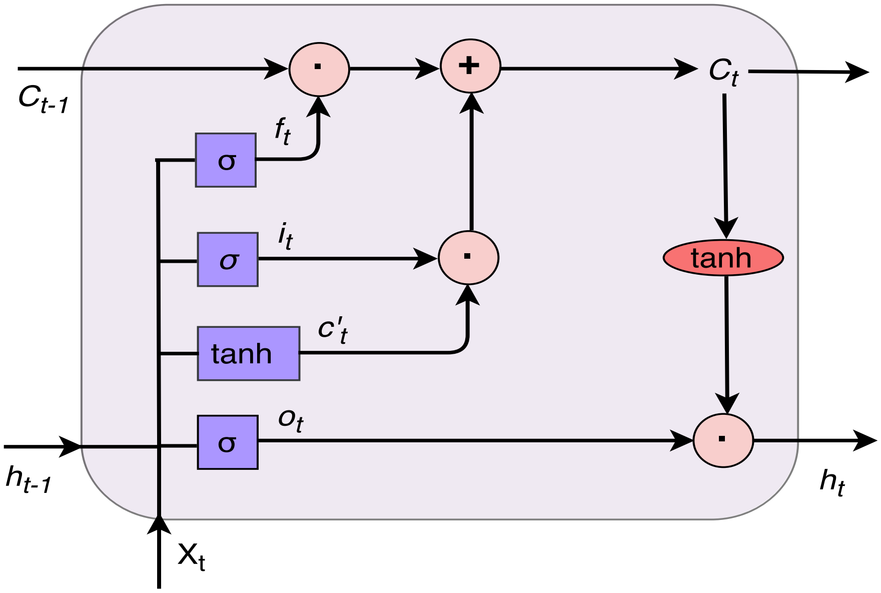


Figure 2: An LSTM cell.

Output from both the hidden layers is then combined and sent to the output layer. The forward hidden sequence ($\overrightarrow{h}$), the backward hidden sequence ($\overleftarrow{h}$), and the output $y_t$ at time step $t$ is given by:

$$\overrightarrow{h}_t = f(W_{x\overrightarrow{h}} x_t + W_{\overrightarrow{h}\overrightarrow{h}} \overrightarrow{h}_{t-1})$$

$$\overleftarrow{h}_t = f(W_{x\overleftarrow{h}} x_t + W_{\overleftarrow{h}\overleftarrow{h}} \overleftarrow{h}_{t+1})$$

$$y_t = f(W_{\overrightarrow{h}y} \overrightarrow{h}_t + W_{\overleftarrow{h}y} \overleftarrow{h}_t)$$

**Attention mechanism** Attention based neural networks are successful in diverse tasks like speech recognition (Chorowski *et al.*, 2015) and language translation (Bahdanau *et al.*, 2015), where the neural network finds it difficult to comprehend the fixed length vector representation of a very long input sequence. We implemented the traditional attention mechanism proposed in (Bahdanau *et al.*, 2015). For an input sequence of length $n$, the importance of each nucleotide position $i$ is accessed by calculating $\alpha_i^j$ for $i = 1, 2, ..., n$ using the formula

$$\alpha_i^j = \frac{exp(e_{ij})}{\sum_{k=1}^{n} exp(e_{ik})}$$

where $e_{ij}$ is the *tanh* activations of the dot products of hidden layer representations ($h_i$), generated by the BLSTM layer, with the attention layer weights. Based on the importance $\alpha_i^j$ of each hidden state $h_i$ generated for input nucleotide $i$, a context vector $c$ is generated as

$$c_j = \sum_{i=1}^{n} \alpha_i^j h_i$$

for predicting the output at time step $j$. At each time step, a new set of attention weights and hidden states are generated. We use the attention weights generated at the $n^{th}$ time step.

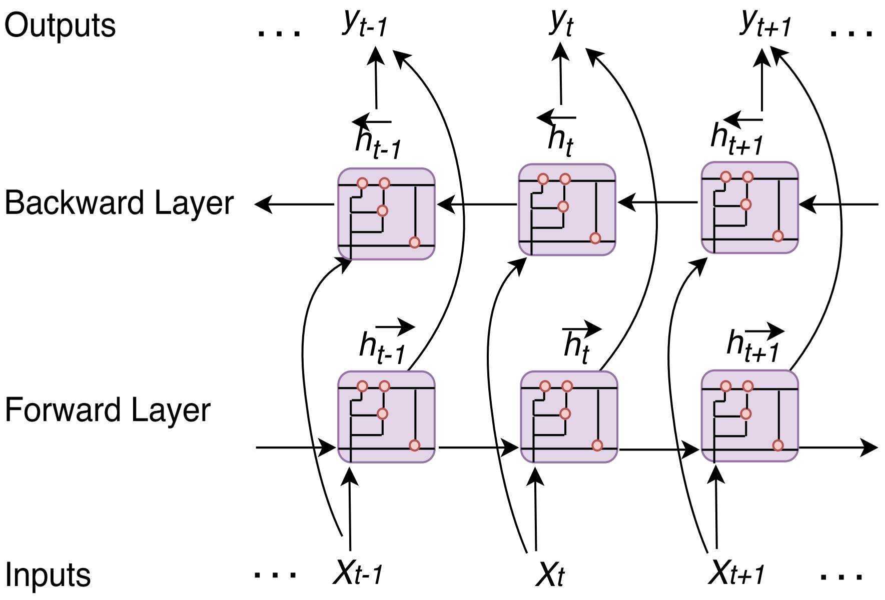


Figure 3: A BLSTM network.

### 2 Experimental Setup

#### 2.1 Efficient implementation of occlusion

Instead of naively iterating through each sequence and then iterating through each window, we prepared the modified sequences for each input sequence beforehand. Thus, for a single input sequence of length $n$, we obtained the modified $n$ sequences corresponding to each occluded window. We concatenated the $n$ modified sequences, resulting in a $n \times n$ matrix corresponding to each input sequence. Finally, upon concatenation of all the modified sequences of the $B$ input sequences in a batch, we obtain a $(B * n) \times n$ matrix which is fed in a single batch to the model, to directly predict the probabilities, resulting in a $(B * n) \times 1$ matrix. This is reshaped to get the required $B \times n$ matrix of *deviation values*.

#### 2.2 Performance comparison of splice junction and junction pair

We compare the performances of various models in the prediction of only a donor junction with flanking regions as well as a junction pair with flanking regions. We observe that all the models perform better in prediction of junction pairs compared to prediction of donor junctions alone.

Table 1: Performance of the various models in donor splice junction and junction pair prediction. Accuracy (Ac), Precision (Pr), Recall (Re) and F1 Score (F1) are computed in percentage.

| Model | Donor prediction | Splice junction prediction |
|---|---|---|
| | $Ac, Pr, Re, F1(\%)$ | $Ac, Pr, Re, F1(\%)$ |
| SpliceVisuL | 92.98, 96.19, 89.52, 92.74 | 98.38, 99.38, 97.38, 98.37 |
| DeepSplice | 90.67, 91.10, 90.15, 90.62 | 92.85, 96.43, 89.03, 92.56 |
| SpliceRover | 88.68, 87.35, 90.61, 88.90 | 90.13, 89.90, 90.41, 90.16 |
| SpliceVec-MLP | 74.19, 71.31, 80.95, 75.82 | 93.98, 96.43, 91.40, 93.78 |

#### 2.3 Data generation

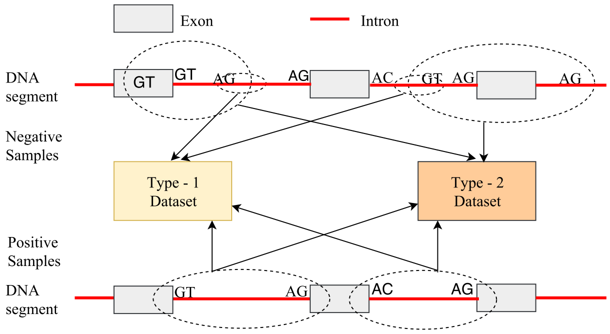


Figure 4: A pictorial representation of the Type-1 and Type-2 dataset. For simplicity of representation, the entire extracted DNA region for the positive samples and the Type-2 negative samples are displayed using the dashed circles. These samples are truncated to 40 nt upstream and downstream regions before feeding into the learning model.

#### 2.4 Variation in length of flanking region

We vary the length of flanking region in the input sequences to find the optimal region that provides best predictive performance of the model. The flanking region is varied from 10 nt to 50 nt in the upstream and downstream sequences of both the donor and acceptor splice junctions. Results shown are for the Type-2 dataset.

The performance of the model improves with increase in the flanking region, displaying the best performance with 40 nt flanking region. All the analysis produced in the paper are performed with a flanking region of 40 nt. The performance of the model with varying flanking region is shown in Table 2.

Table 2: Performance of the model with variation in flanking region

| Flanking region | Performance measures |
|---|---|
| | $Ac, Pr, Re, F1(\%)$ |
| 50 nt | 97.97, 98.91, 97.00, 97.95 |
| 40 nt | **98.38, 99.38, 97.38, 98.37** |
| 30 nt | 97.18, 99.03, 95.29, 97.13 |
| 20 nt | 96.45, 97.26, 95.60, 96.42 |
| 10 nt | 95.06, 95.71, 94.35, 95.03 |

#### 2.5 Variation in model architecture

Table 3 shows the variation in the performance of the model with variation in the number of hidden layers (HL). Results are shown for Type-2 dataset. We show the performance with both LSTM and BLSTM as the units of the hidden layer. We see that the model is invariant to the number of hidden layers as well as the type of units in the hidden layer. All the analysis produced in the paper are performed with a model having one hidden layer of BLSTM units.

#### 2.6 Hyperparameters of the various baselines

Discussed below are the hyperparameters tuned for various baselines.

- Vanilla LSTM: We consider a batch size of 128, dropout rate as 0.5, recurrent dropout rate as 0.2, and number of epochs as 50.

Table 3: Performance of the model with variation in architecture

| Model | Performance measures |
|---|---|
| | $Ac, Pr, Re, F1(\%)$ |
| LSTM with 1 HL | 97.68, 98.88, 96.45, 97.65 |
| LSTM with 2 HL | 98.49, 99.10, 97.88, 98.48 |
| BLSTM with 1 HL | 98.38, 99.38, 97.38, 98.37 |
| BLSTM with 2 HL | 98.24, 99.66, 96.81, 98.21 |

- LSTM with attention: We consider a batch size of 128, dropout rate as 0.5, recurrent dropout rate as 0.2, and number of epochs as 10.
- SpliceVec-MLP (Dutta *et al.*, 2018): We consider one hidden layer with 2500 hidden nodes. We used a batch size of 128 and the learning rate of the Adam optimizer as 0.001.
- DeepSplice (Zhang *et al.*, 2016): We consider a batch size of 160, dropout keep probability as 0.8 and the Adam optimizer learning rate as 0.001.
- SpliceRover (Zuallaert *et al.*, 2018): We consider a batch size of 64, learning rate as 0.05 with decay rate of 0.5 every 5 steps and Stochastic Gradient Descent with Nesterov momentum 0.9 as the update strategy. We consider only one convolutional and one max-pooling layer as this architecture provided the best performance.

#### 2.7 Input formation for SpliceVec-MLP

This model extracts the complete intronic sequence, along with the flanking upstream and downstream region, converts it into an N-dimensional feature representation, and then feeds it into the learning model. In Type-1 dataset, the negative splice junctions are generated by extracting a continuous sequence of 164 nt from the middle of each intron. Unlike Type-2 dataset, Type-1 dataset has a fixed length negative input samples.

Hence, to access the performance of SpliceVec-MLP on Type-1 dataset, we have truncated the positive splice junction sequences to 40 nt upstream and downstream region of both the donor and acceptor splice junctions. The 82 nt sequences obtained from both the splice junctions of a junction pair were concatenated to obtain a 164 nt sequence representing a true splice junction.

#### 2.8 Existing consensus for analyzing visualizations

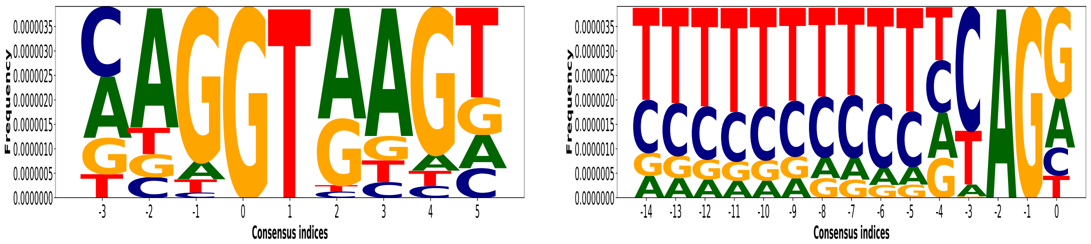


(a) (b)

Figure 5: Existing consensus in the canonical (a) donor and (b) acceptor splice junctions of the dataset. The consensus comprises the extended donor site consensus 9-mer $[AC]AGGTRAGT$ and the extended acceptor site consensus 15-mer $Y_{10}NCAGG$.

### 3 Results

#### 3.1 Prediction performance

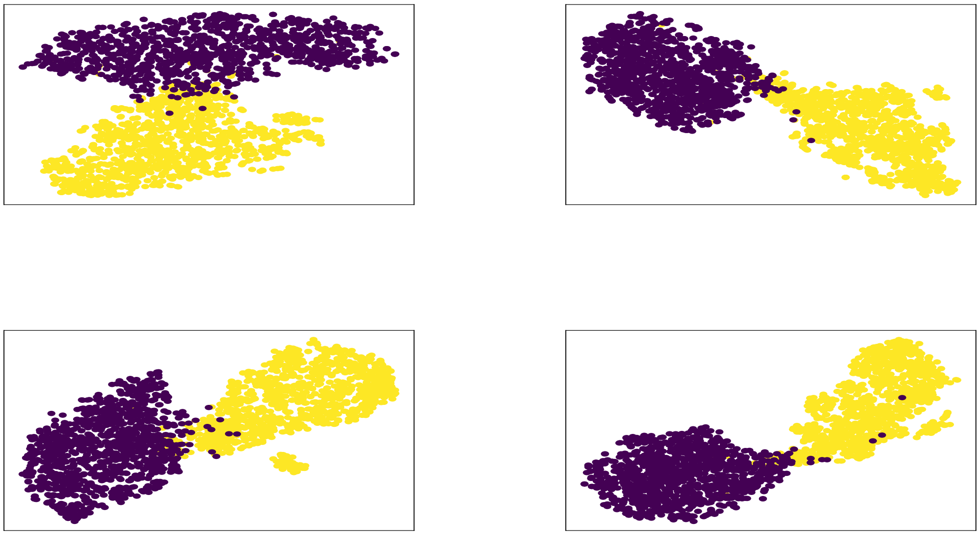

Figure 6: t-SNE plots of 4 different sets of 1000 true and 1000 decoy splice junctions. Points in blue represent true whereas points in yellow represent decoy splice junctions. We have plotted 1000 sequences considering the execution time required for larger sample sizes.

#### 3.2 Deviation values per position

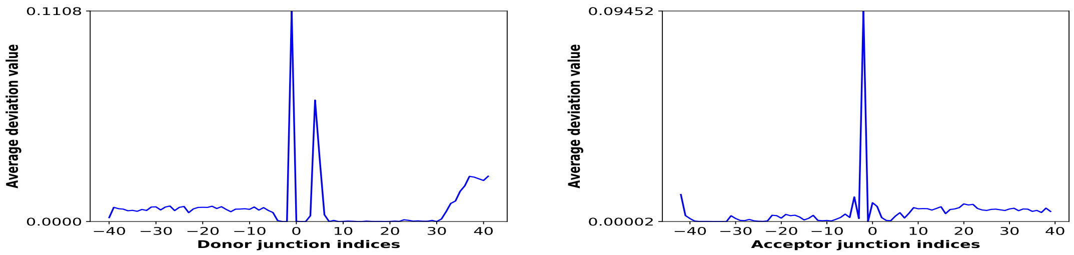


Figure 7: Average deviation value per position based on attention weights of canonical sequences.

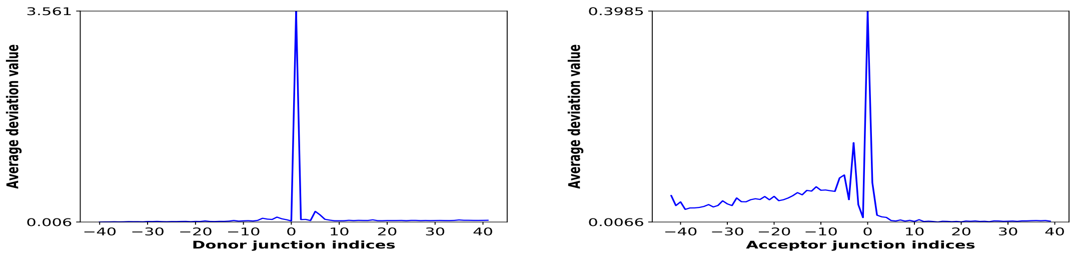


Figure 8: Average deviation value per position based on smooth gradients of canonical sequences.

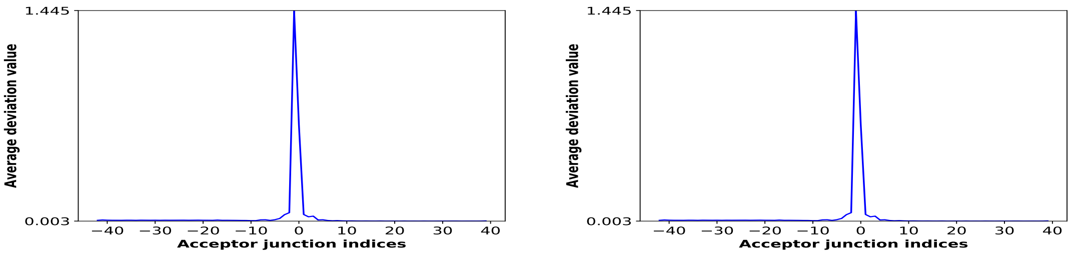


Figure 9: Average deviation value per position based on integrated gradients of canonical sequences.

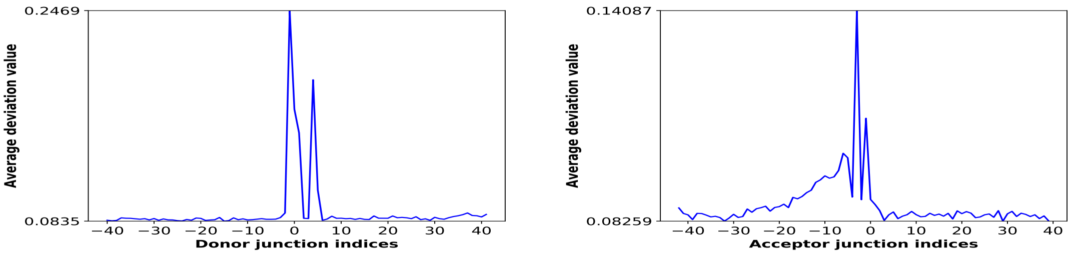


Figure 10: Average deviation value per position based on omission of canonical sequences.

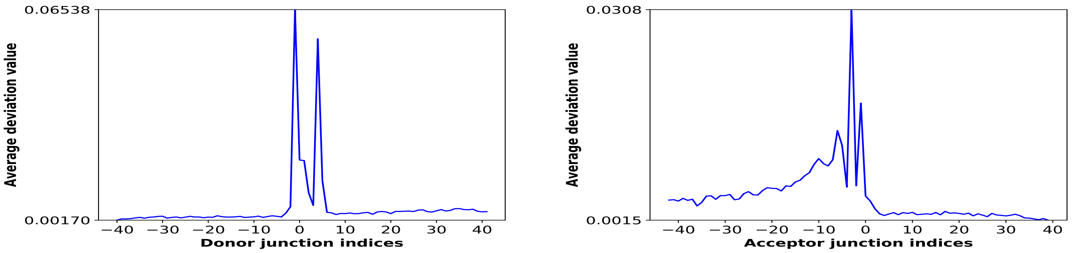


Figure 11: Average deviation value per position based on occlusion-1 of canonical sequences.

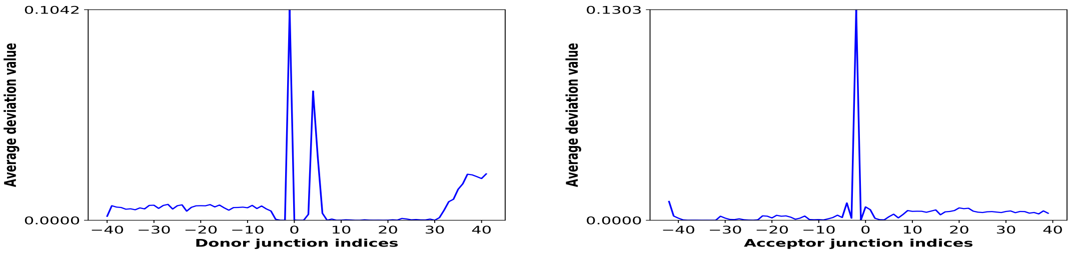


Figure 12: Average deviation value per position based on attention weights of non-canonical sequences.

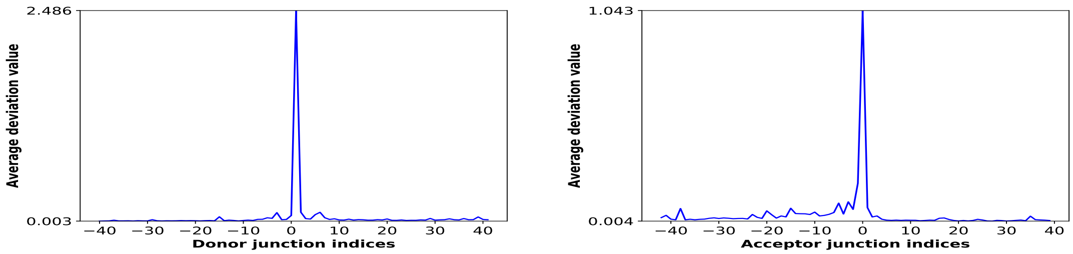


Figure 13: Average deviation value per position based on smooth gradients of non-canonical sequences.

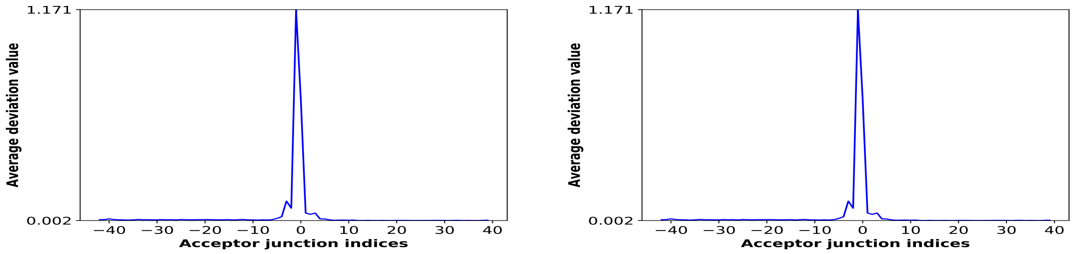


Figure 14: Average deviation value per position based on integrated gradients of non-canonical sequences.

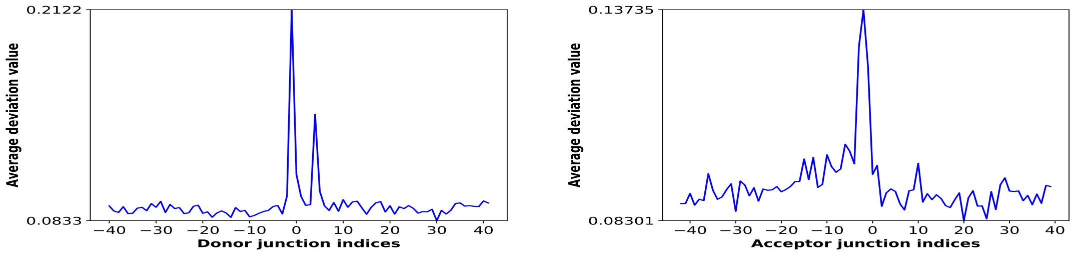


Figure 15: Average deviation value per position based on omission of non-canonical sequences.

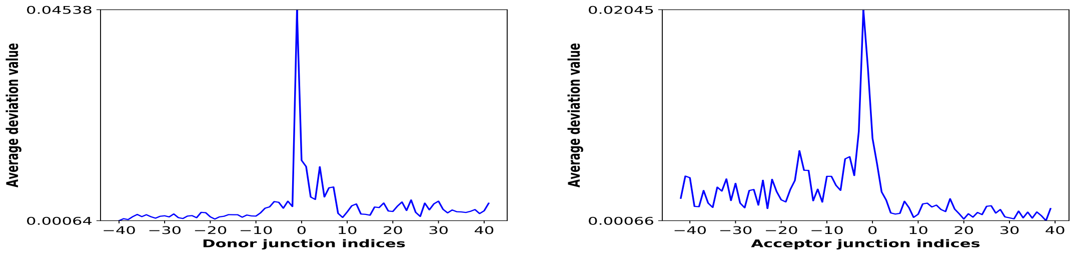


Figure 16: Average deviation value per position based on occlusion-1 of non-canonical sequences.

#### 3.3 Deviation values per position per nucleotide

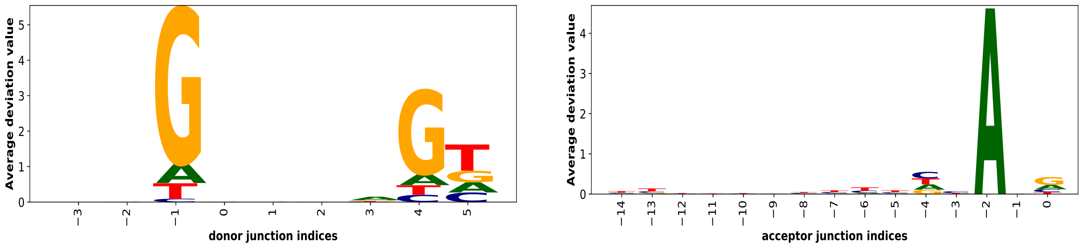


Figure 17: Average deviation value per position per nucleotide based on attention weights of canonical sequences.

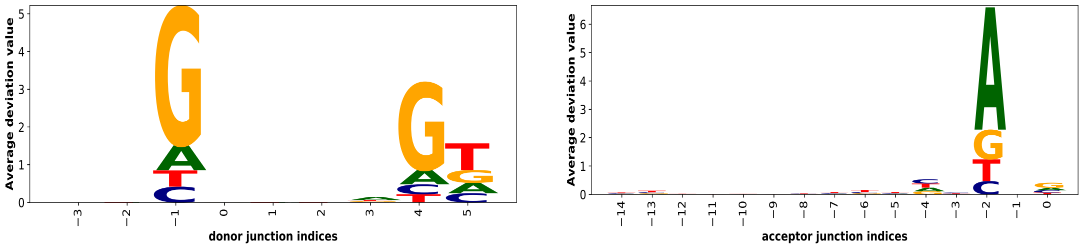


Figure 18: Average deviation value per position per nucleotide based on attention weights of non-canonical sequences.

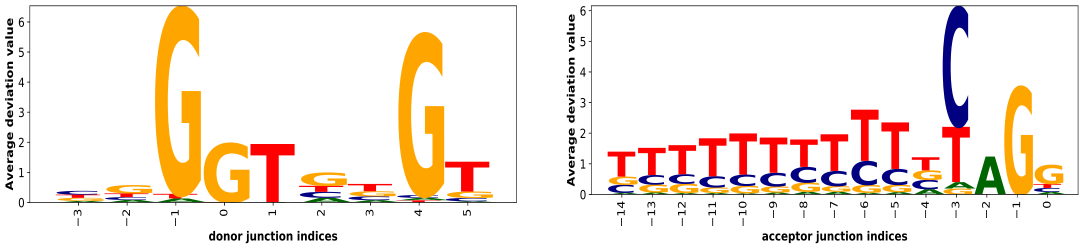


Figure 19: Average deviation value per position per nucleotide based on occlusion-1 of canonical sequences.

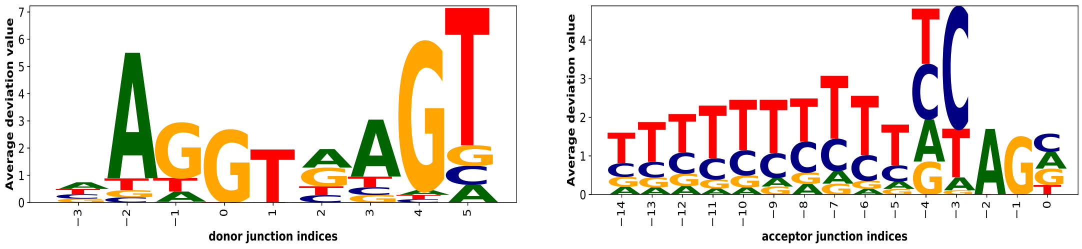


Figure 20: Average deviation value per position per nucleotide based on occlusion-3 of canonical sequences.

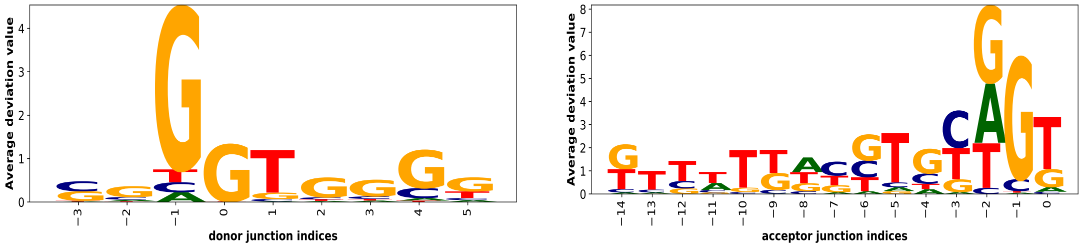


Figure 21: Average deviation value per position per nucleotide based on occlusion-1 of non-canonical sequences.

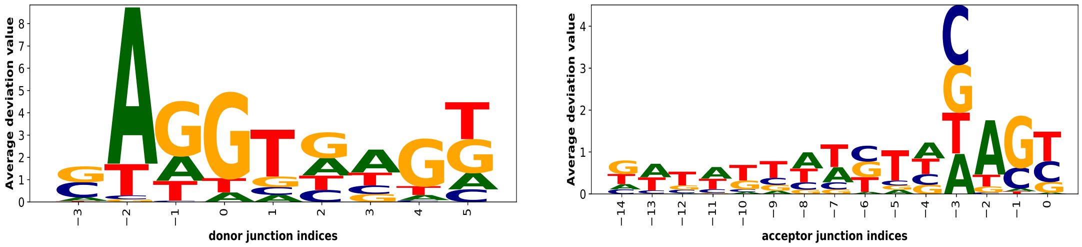


Figure 22: Average deviation value per position per nucleotide based on occlusion-3 of non-canonical sequences.

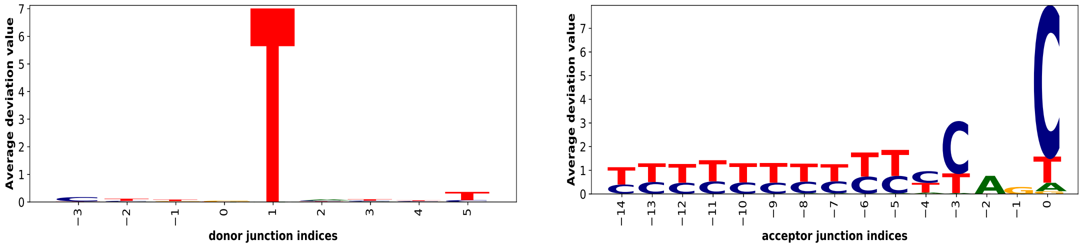


Figure 23: Average deviation value per position per nucleotide based on smooth gradients of canonical sequences.

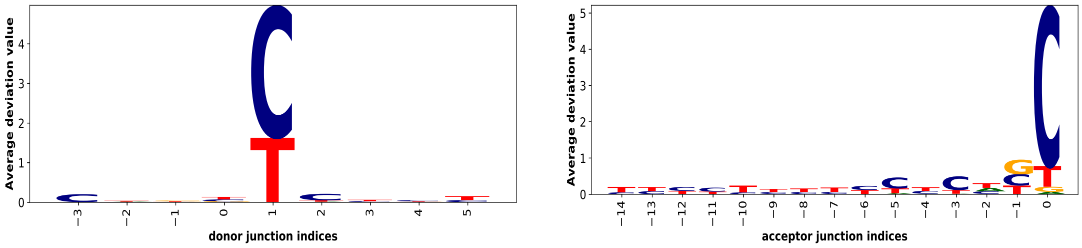


Figure 24: Average deviation value per position per nucleotide based on smooth gradients of non-canonical sequences.

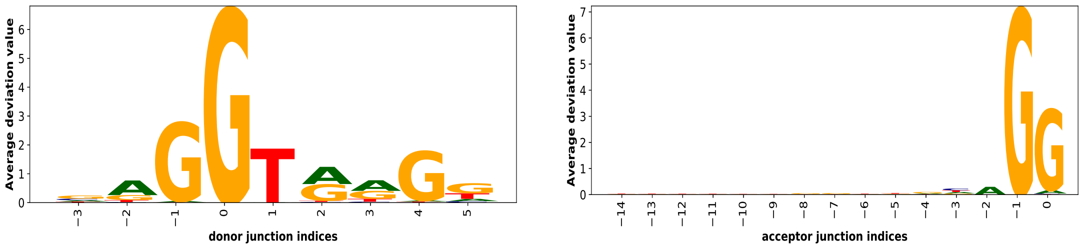


Figure 25: Average deviation value per position per nucleotide based on integrated gradients of canonical sequences.

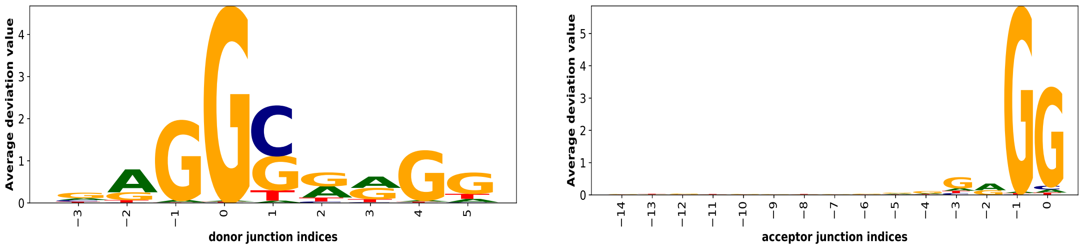


Figure 26: Average deviation value per position per nucleotide based on integrated gradients of non-canonical sequences.

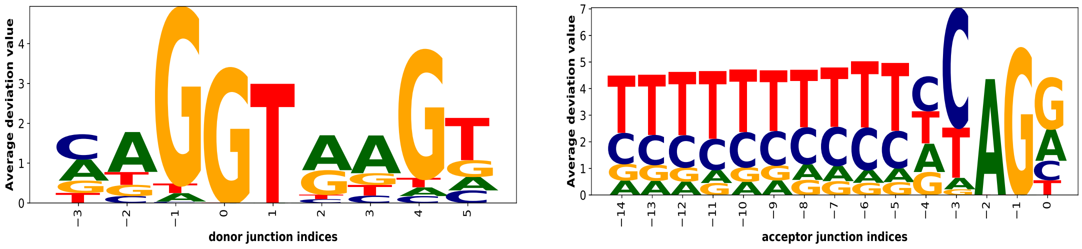


Figure 27: Average deviation value per position per nucleotide based on omission scores of canonical sequences.

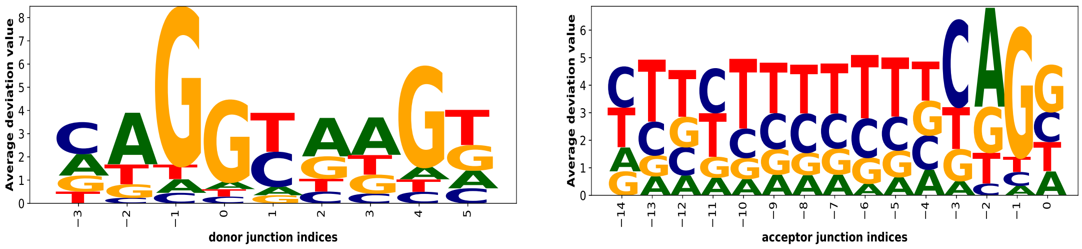


Figure 28: Average deviation value per position per nucleotide based on omission scores of non-canonical sequences.

#### 3.4 Deviation values per position for branchpoint pattern ‘CTRAY’ in the region -40 to -15

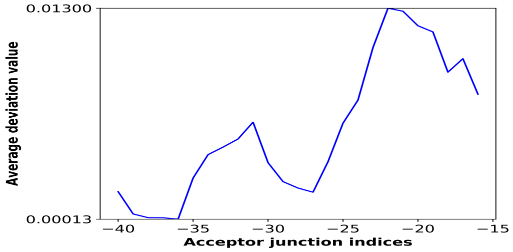


(a)

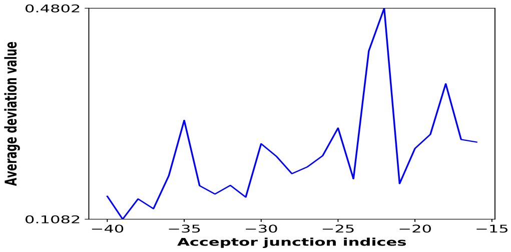


(b)

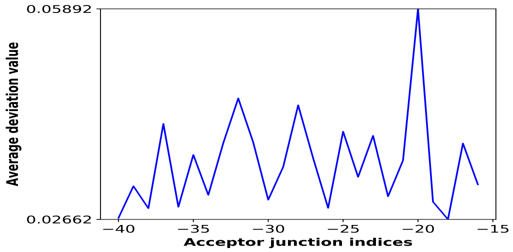


(c)

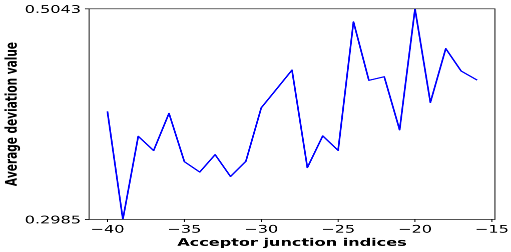


(d)

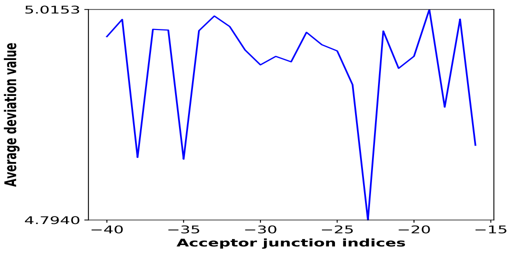


(e)

Figure 29: Average deviation value per position for branchpoint pattern ‘CTRAY’ based on (a) attention (b) smooth gradients (c) integrated gradients (d) omission and (e) occlusion-3 of canonical splice junctions respectively.

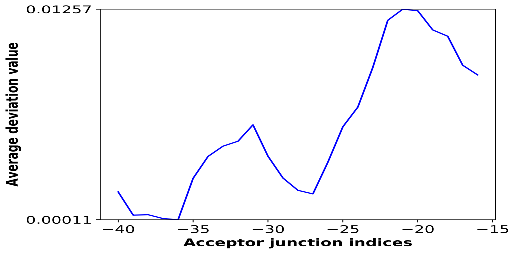


(a)

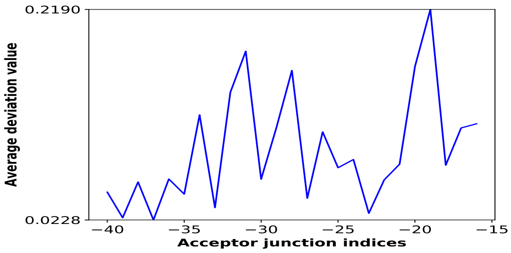


(b)

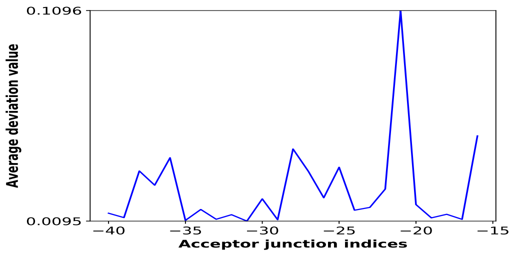


(c)

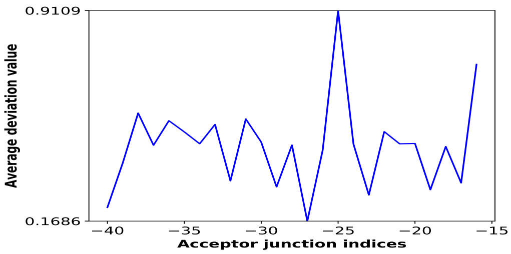


(d)

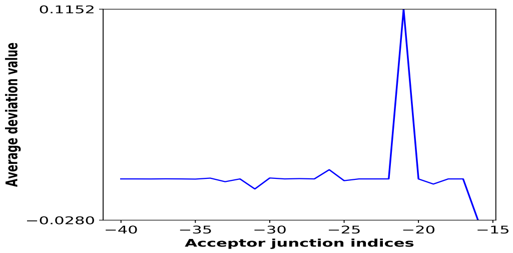


(e)

Figure 30: Average deviation value per position for branchpoint pattern 'CTRAY' based on (a) attention (b) smooth gradients (c) integrated gradients (d) omission and (e) occlusion-3 of non-canonical splice junctions respectively.
